# Pre- and postsynaptic nanostructures increase in size and complexity after LTP induction

**DOI:** 10.1101/2023.02.09.527812

**Authors:** Valérie Clavet-Fournier, ChungKu Lee, Waja Wegner, Nils Brose, JeongSeop Rhee, Katrin I. Willig

## Abstract

Synapses, specialized contact sites between neurons, are the fundamental elements of neuronal information transfer. Synaptic plasticity is related to changes in synaptic morphology and the number of neurotransmitter receptors, and thought to underlie learning and memory. However, it is not clear how these structural and functional changes are connected. We utilized time-lapse super-resolution STED microscopy to visualize structural changes of the synaptic nano-organization of the postsynaptic scaffolding protein PSD95, the presynaptic scaffolding protein Bassoon, and the GluA2 subunit of AMPA receptors by chemically induced long-term potentiation (cLTP) at the level of single synapses. We found that the nano-organization of all three proteins undergoes an increase in complexity and size after cLTP induction. The increase was largely synchronous, peaking at ∼60 min after stimulation. Therefore, both the size and complexity of single pre- and post-synaptic nanostructures serve as substrates for adjusting and determining synaptic strength.

**Highlights:**

- Time-lapse super-resolution images the structural changes of the PSD95 nano-organization after Cltp
- cLTP-induced growth of the PSD95 nano-organization is less than spine head growth and peaks at 60 min, i.e. much slower than the increase in spine volume.
- Most PSD95 nanostructures increase in complexity upon cLTP.
- Synchronous growth - Nanostructures of pre- and postsynaptic scaffolding proteins and AMPA receptors increase simultaneously and equally strong upon cLTP.
- GluA2-containing synaptic AMPA receptors form nanoclusters that increase in size and slightly in number upon cLTP and form subdomains on PSD95.
- Bassoon forms complex structures similar to the PSD95 nano-organization.

## Introduction

Chemical synapses, which consist of presynaptic and postsynaptic compartments as well as a synaptic cleft, employ a multitude of exquisitely arranged and coordinated proteins to convert electrical into chemical signals, and vice versa. The rate and efficacy of synaptic transmission varies among synapses, and is thus unique to each synapse. This diversity is in large part due to differences in release probability of synaptic vesicles at the presynapse and the heterogenous distribution of neurotransmitter receptors at the postsynapse. However, it is still debated whether and how synaptic strength correlates with changes in the structural organization of synapses and their protein components. Decades ago it was shown by electron microscopy that the postsynaptic density (PSD), a proteinaceous specialisation of excitatory postsynapses is not always continuous but often disrupted or perforated. Such perforations occur mainly on larger postsynaptic spines, specialized dendritic protrusions of excitatory glutamatergic postsynapses, and can be of various, complex shape^1^. The proportion of perforated synapses and their size were shown to increase with development or the induction of long-term potentiation (LTP)^2,3^. Recently, such rapid structural changes of the PSD and non-synaptic compartments were quantified in detail after induction of LTP in single spines^2^. While synaptic perforations are too small to be visible by conventional light microscopy, the application of super-resolution techniques enables their visualization in living cells, even in the intact brain of a living mouse. We and others showed that the postsynaptic density protein PSD95, a scaffolding protein highly enriched in excitatory postsynapses, is also not always assembled in a continuous structure but in nanoclusters or more complex shapes^3–5^; these assemblies change their morphology at baseline *in vivo* at the minute to hour time-scale^5^ and increase in size upon enhanced activity in an enriched environment^6^. While spines increase rapidly in volume after LTP induction, PSD95 assemblies increase in size much more slowly, over a few hours^7,8^. The recent discovery that components of the presynaptic active zone line up with proteins of the postsynaptic density and glutamate receptors in so-called nanocolumns^9,10^ intensified the investigation of the structural nano-organization of the synapse and its functional significance.

However, the corresponding studies have several caveats. First, studies using conventional two-photon microscopy are limited in resolution and cannot resolve the PSD95 nanostructure. Second, most studies using super-resolution microscopy to address the synaptic nano-organization of PSD95 assessed only the size and/or number of clusters, so called nano-clusters, nano-domains, or nano-modules^3,4,9,10^. However, with *in vivo* STED microscopy of an adult knock-in mouse expressing PSD95-eGFP we found that PSD95 is nano-organized in various shapes, such as perforated, ring-like, or more complex structures^5^. This is consistent with results from electron microscopy^2^. Thus, a simple cluster analysis alone misses key features required for our understanding of the functional role of the complex synaptic nanostructure. Third, studies of activity-dependent changes in the synaptic nano-organization usually compare different datasets collected at different time points after stimulation and are therefore indirect measurements; this holds true for super-resolution imaging^9,11^ and electron microscopy^2^.

Here, we overcome these limitations by employing time-lapse super-resolution STED microscopy of endogenous PSD95 in living organotypic hippocampal slices. We investigated the plasticity of spines and PSD95 nano-organization upon NMDA dependent chemical LTP (cLTP) and assessed changes in size and nanostructure similar to a recent electron microscopic study^2^. PSD95 nano-organizations increased in complexity and size after cLTP induction, with the restructuring of PSD95 assemblies occurring more slowly than the rapid growth of the spine head. Furthermore, we tested whether arrangements of presynaptic scaffolds and glutamate receptors are equally complex in nanostructure and dynamics. We found a strong correlation between the nano-organization of presynaptic Bassoon and postsynaptic PSD95; Bassoon clusters are similarly complex in structure and increase in size by a similar factor upon cLTP induction. In contrast, AMPA receptor (AMPAr) nanoclusters containing the GluA2 subunit do not show a complex structure. cLTP induction resulted in an increase of synaptic AMPAr nanocluster size and number, contradicting a pure modular composition, which was previously suggested^11^. That is, the change in the AMPAr-induced excitatory postsynaptic current (EPSC) size was more sensitive and rapid than the dynamics of AMPAr nanoclusters, which was similar to the slow change in PSD95 assemblies.

## Results

### Time-lapse STED imaging of endogenous PSD95 in organotypic hippocampal slices

To assess the morphology and nano-plasticity of PSD95, we used a home-built STED microscope^12,13^ to image PSD95 in live organotypic hippocampal slices. PSD95 was highlighted with a transcriptionally regulated recombinant antibody-like protein termed FingR, labeling endogenous PSD95^6,14^. Additionally, we expressed a myristoylation (myr) motif to tag the neuronal membrane and reveal spine morphology. To detect the fluorescence of the Citrine tagged PSD95.FingR in a broad spectral range, we used the reversibly switchable fluorescent protein (RSFP) rsEGFP2 for the membrane^13^. rsEGFP2 emits fluorescence very close to Citrine (Figure 1A) and therefore both fluorescent proteins (FP) can be excited and detected in the same broad spectral window. Fluorescence of rsEGFP2 was recorded sequentially in time by switching rsEGFP2 to the on state with 405 nm light (Figure 1B). For recording the pure PSD95 image rsEGFP2 was in the off state and thus not fluorescent (Figure S1). This off-switching was performed by the blue excitation laser while recording the Citrine fluorescence. In this manner we revealed the PSD95 nano-organization in super-resolution; spine morphology was recorded in confocal mode only, as this was sufficient to determine the size of the spine heads (Figure 1C). Both constructs were expressed via transduction of recombinant adeno-associated viral particles (AAV) into the CA1 region of the hippocampus. Recording images at different time points revealed the structural changes of the PSD95 nano-organization (Figure 1D). For further analysis we encircled the spine head and PSD95 assemblies to assess their size (Figure 1E).

**Figure 1:**
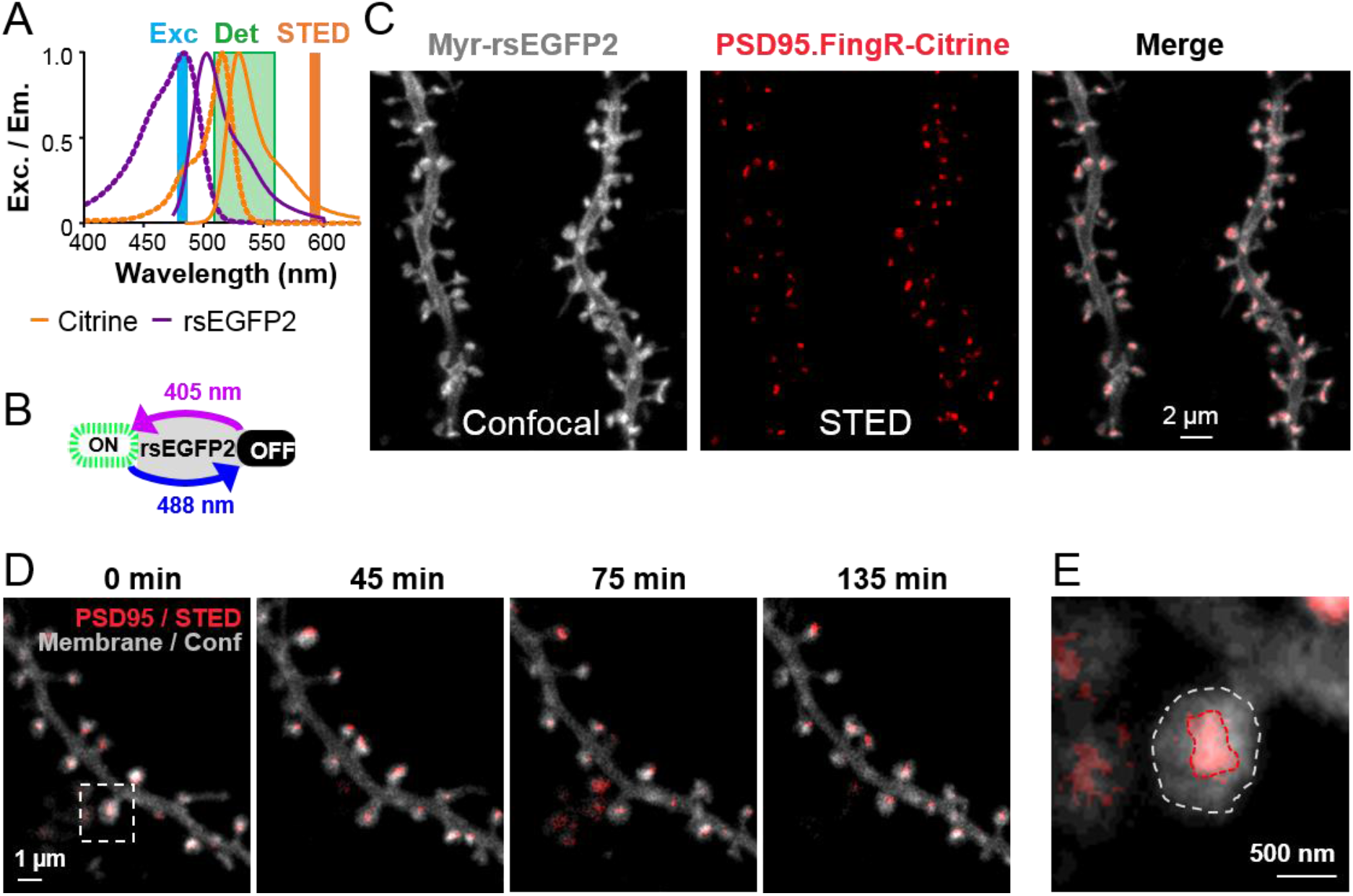
Super-resolution microscopy of endogenous PSD95 in hippocampal organotypic slices. **(A, B)** Dual-label schema by sequential read out: The FPs Citrine and rsEGFP2 are excited with blue light (Exc) and detected between 510–560 nm (Det); stimulated emission depletion (STED) is performed at 595 nm (A). The reversibly switchable FP rsEGFP2 is switched on demand to the on state with UV light at 405 nm and to the off state with blue light at 488 nm. **(C)** Hippocampal neurons express a myristoylation tag (myr) and a dendrite targeting sequence (LDLR, cf. Methods) fused to rsEGFP2 to label the dendritic membrane and an antibody-like tag (PSD95.FingR) fused to Citrine to label endogenous PSD95. Super-resolution STED microscopy of PSD95 (red) and confocal imaging of the neuronal membrane (white). **(D)** Time-lapse imaging of PSD95 and spine morphology for > 2 hours. **(E)** Magnification of boxed area in (D); encircling of the PSD95 assembly and spine head for size analysis. (C, D) Images are smoothed and maximum intensity projection.

### PSD95 assembly size and synaptic strength increase upon cLTP

To further explore structural and functional synaptic plasticity, we induced cLTP by adding 200 μM glycine and 20 μM bicuculline to ACSF without Mg^2+^ ions (Figure 2A). Before cLTP induction, the hippocampal organotypic slices were kept in ACSF solution containing APV to block basal neuronal activity induced by NMDA receptors for 20 min. To assess the functional changes of single synapses induced by cLTP we recorded spontaneous excitatory events by whole cell voltage clamp recordings in pyramidal neurons of CA1^15–17^, with the rationale that the amplitude and kinetics of corresponding miniature excitatory postsynaptic currents (mEPSC) reflect, in part, functional and structural changes occurring at synapses. mEPSCs were measured for up to one hour after cLTP induction, while all solutions contained TTX to avoid effects caused by propagated action potentials. mEPSC amplitudes increased immediately after cLTP induction protocol by a factor of 1.28 ± 0.05 and enhanced currents were maintained for at least 60 min; without stimulation mEPSC amplitudes did not increase (0.93 ± 0.03; control) (Figure 2B).

**Figure 2:**
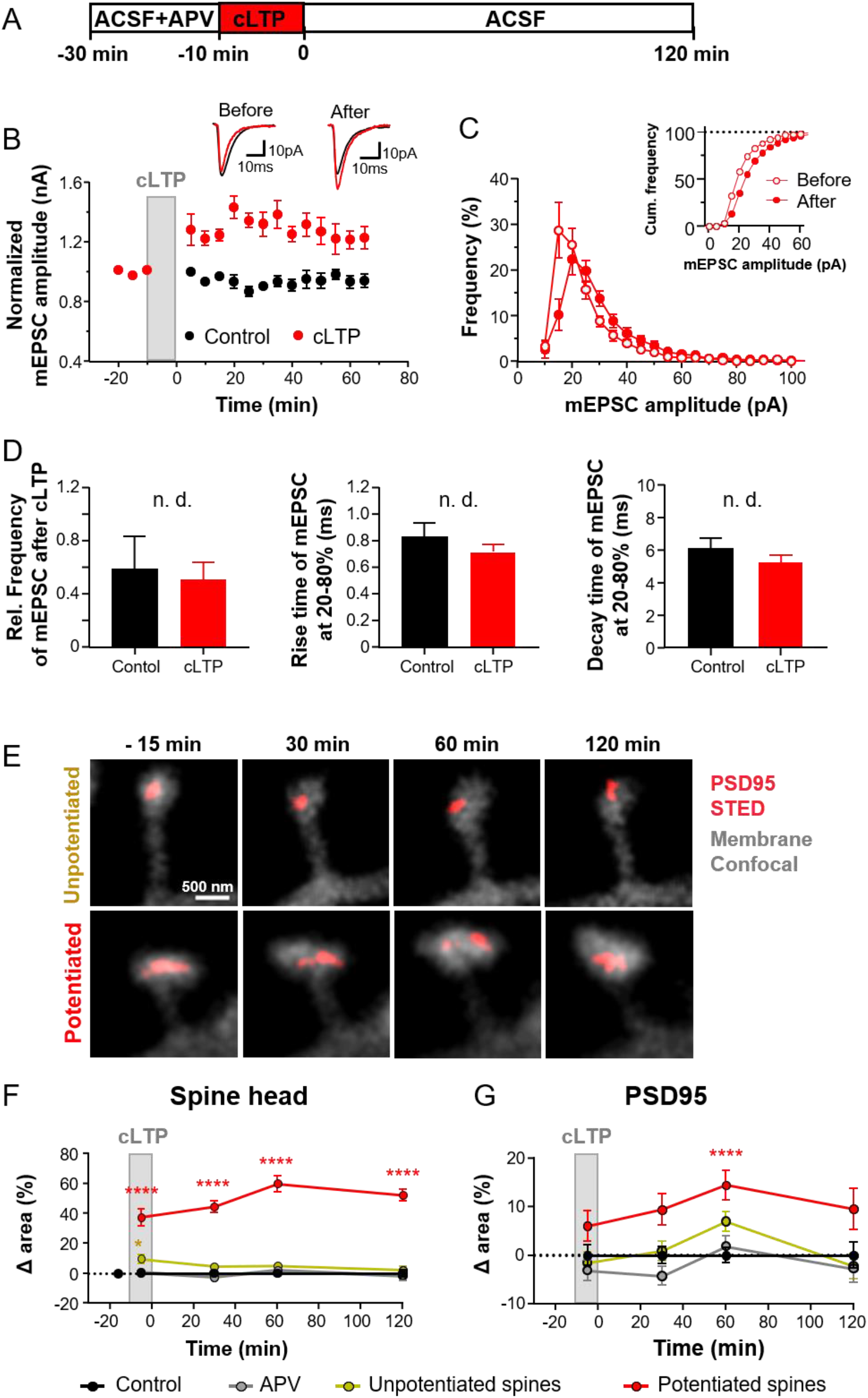
Increase in synaptic strength and PSD95 assembly size after cLTP induction. **(A)** Time-line of the experiment. Chemical LTP (cLTP) is induced by ACSF containing zero Mg^2+^, 200 μM glycine, 20 μM bicuculline. **(B–D)** Whole cell voltage clamp recording of mEPSC in organotypic hippocampal slices. (B) Representative mEPSC traces (top) of CA1 pyramidal neurons before (−15min) and after (65 min) cLTP and averaged normalized current ± SEM (bottom); control without chemical stimulation (black) and cLTP (red). (C) Frequency distribution of mEPSC amplitude and cumulative frequency before (open circle, -15 min) and after cLTP induction (closed circle, 65 min). (D) Normalized frequencies of mEPSC (left), rise time (middle) and decay times (right) for 20–80 % of mEPSC. Bars represent average ± SEM with (red) and without (black) induction of cLTP; no significant difference between cLTP and control (unpaired t-test) **(E)** Representative images of potentiated and unpotentiated spines before and after cLTP induction. **(F)** Mean ± SEM of changes in spine head area of potentiated and unpotentiated spines after cLTP relative to control. Control conditions were continuously kept in ACSF (control) and spines at blocked activity measured in ACSF containing APV (APV). Changes were compared to control (Kruskal-Wallis with Dunn’s multiple comparisons test; ****p < 0.0001). **(G)** Mean ± SEM for PSD95 area of the spines analysed in (F) (Kruskal-Wallis with Dunn’s multiple comparisons test; 60 min: ****p < 0.0001). (B–D) Number of recorded cells: cLTP: 9; control: 5. (F, G) Number of analysed spines at -5, 30, 60, 120 min: Control: 123, 224, 223, 101; APV:, 153, 229, 201 84; unpotentiated: 66, 202, 196, 130; potentiated: 46, 106, 111, 65. Number of analysed PSD95 assemblies at the same time points: Control: 118, 220, 211, 92; APV: 153 220 165 69; unpotentiated: 68, 201, 195, 124; potentiated: 45, 97, 105, 56. Number of hippocampal slices respectively cells: Control: 10; APV: 13; unpotentiated: 19; potentiated: 19.

To determine whether cLTP is a phenomenon that affects only a limited number or specific synapses, we further analysed the frequency distribution of mEPSC amplitudes. After cLTP, the distribution of mEPSC amplitudes shifted to the right and widened slightly (Figure 2C). A plot of the normalized frequencies reveals a very similar shape of mEPSC amplitudes before and after cLTP induction (Figure S2B) which indicates that the vast majority of synapses increase in strength by a similar factor. In contrast, a widening alone or, in the extreme case, splitting into two peaks would indicate an increase in synaptic strength solely in a subset of synapses.

The relative changes in mEPSCs frequency did not show any difference between the control (0.52 ± 0.13) and cLTP-induced groups (0.58 ± 0.26) (Figure 2D, left). This indicates that the increase in mEPSC amplitude is not due to an increase in pre-synaptic vesicle release probability, but rather to changes in the number of receptors at the post-synapse. To assess the kinetics of the mEPSC, we analysed the 20–80 % rise time and the decay time of mEPSCs before and after cLTP induction. The rise time was not significantly different between cLTP (0.83 ± 0.1 ms) and control (0.71 ± 0.05 ms) groups (Figure 2D, middle). Likewise, no difference in the decay time was found between cLTP (5.3 ± 0.4 ms) and control groups (6.2 ± 0.6 ms) (Figure 2D, right).

To investigate structural synaptic plasticity after cLTP induction we performed time-lapse STED imaging. After the acquisition of a baseline image, cLTP was induced for 10 min before perfusion was switched to standard ACSF. Image stacks of the PSD95 labeling and spine membrane were recorded about 5 min before cLTP (t = -15 min), at 5 min during cLTP induction (t = -5 min) and 30, 60, and 120 min after cLTP induction. Spines that were within the z-stack at all time points of the measurement series were analyzed for spine head and PSD95 assembly size. Spines that underwent a persistent enlargement of ≥15 % at 60 min and 120 min after cLTP induction were considered potentiated spines. All other spine heads, i.e. with <15 % enlargement, were regarded unpotentiated. Figure 2E shows a representative example of a potentiated and unpotentiated spine. On average, ∼40 % of the spine heads showed a persistent enlargement of at least 15 % at 60 min and 120 min following cLTP induction (111 of 279 spines in 19 cells). Dendrites with less than 20 % of enlarged spines were discarded. As was shown before^7,8^, the heads of potentiated spines increased significantly already during the cLTP induction (Figure 2F) by 37 ± 5 % compared to controls. This enlargement was persistent and increased even further up to 60 ± 5 % after 60 min and 52 ± 4 % after 120 min. Unpotentiated spine heads, in turn, showed only a small but significant increase of 10 ± 3 % during cLTP induction compared to control, which was not sustained for the rest of the time course. PSD95 assemblies on potentiated spines increased slightly, but not significantly, already during cLTP induction. The increase continued over time and was significant at 60 min (Figure 2G); however, with 14 ± 3 % this increase was smaller than the increase in spine head size. Such a delayed increase of PSD95 cluster size vs. spine head size had also been shown in a previous study^7^. All changes were assessed in relation to changes of a baseline control condition, which was measured in ACSF without stimulation. Blocking of NMDA receptors with APV during the whole-time course prevented changes in spine head or PSD95 assembly size (Figure 2F, G).

### Restructuring of the PSD95 nano-pattern after cLTP induction

In large spine heads, PSD95 is often organized in very complex structures that include perforations, U-shapes, or more complex shapes; it appears in single or in multiple patches (Figure 3A). We have observed a similar PSD95 nano-pattern in a knock-in reporter mouse line^18^ and also *in vivo* with the same FingR.PSD95 construct that we used in the present study^6^. Similar shapes were also observed with electron microscopy of post-synaptic densities^19^. We categorized the PSD95 nanoarchitecture as follows^19^: (1) Simple PSD95 assemblies without a perforation were assigned a macular shape; (2) assemblies with a hole, U-shape or more complex shape were assigned a perforated shape; (3) two separated PSD95 spots were assigned segmented 2; (4) three or more separated PSD95 spots were classified as segmented ≥3. Segmented spots were not further classified for their shape. We performed this classification for spines during and after cLTP induction and for unstimulated controls as described above. In the control, the proportion of the different shapes was mainly conserved over the time course (Figure 3B). About 93 % of the PSD95 assemblies were of macular shape before cLTP at -15 min, 90 % at 60 min, and 91 % at 120 min. Similarly, the proportions of perforated, segmented 2, and segmented ≥3 PSD95 assemblies was conserved in control. However, in the case of potentiated spines, the proportion of macular PSD95 assemblies was 91 % before cLTP induction, but decreased to 86 % at 30 min, 78 % at 60 min, and 73 % at 120 min after cLTP (Figure 3B). This reduction of macular shapes was accompanied by an increase of segmented 2 and perforated PSD95 assemblies. For instance, the number of segmented 2 was 7 % before cLTP and increased up to 12 % at 30 min, 16 % at 60 min and 17 % at 120 min. Only 2 % of the PSD95 assemblies were perforated before cLTP. This value increased to 4 % after 60 min and 8 % at 120 min after stimulation (Figure 3B). Altogether, this time-course shows that cLTP induction affects the PSD95 nano-pattern where PSD95 is reorganized into more complex, perforated, and segmented shapes after cLTP induction.

**Figure 3:**
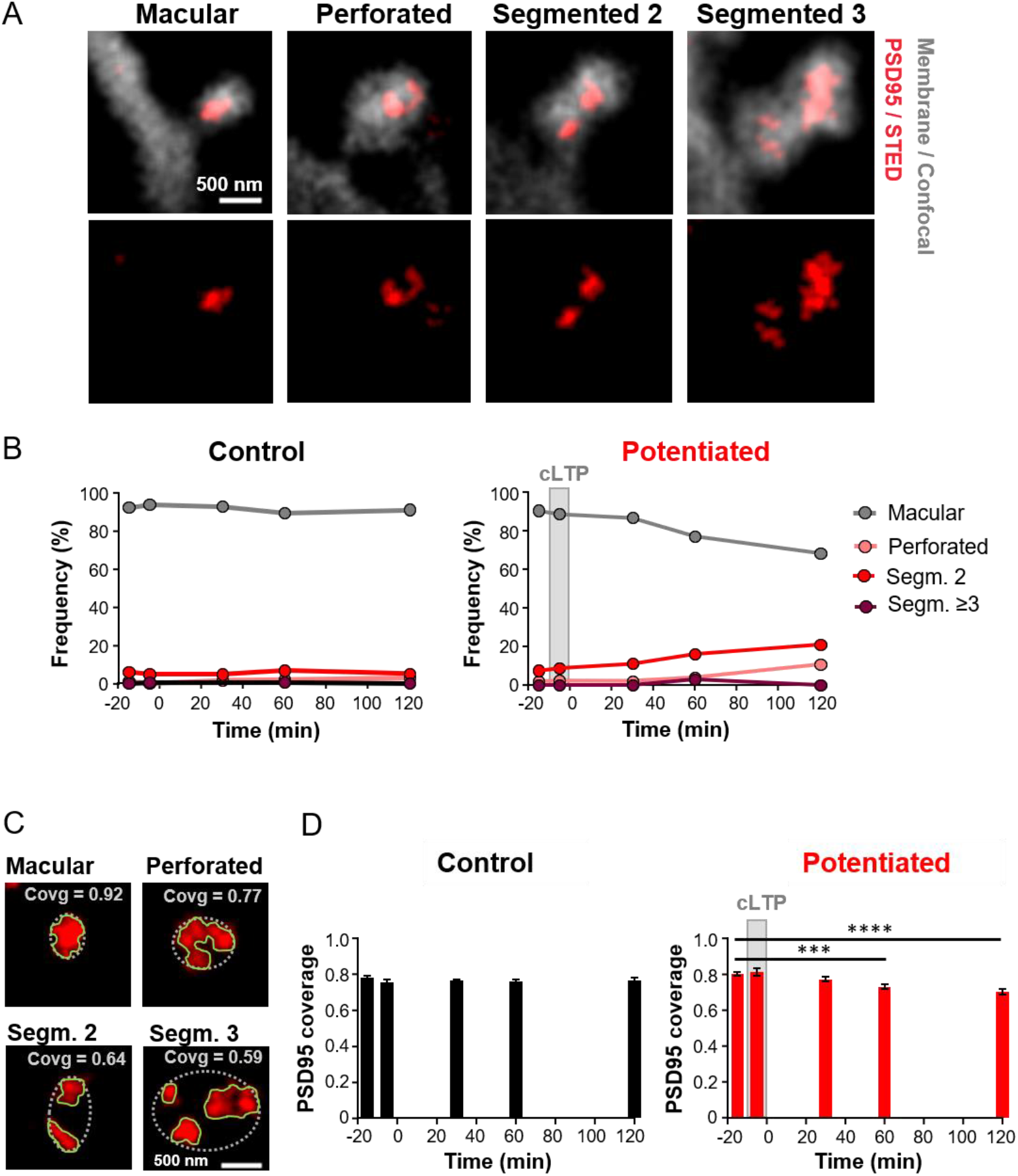
Reorganization of the PSD95 nano-pattern after cLTP induction. **(A)** STED images of PSD95 (red) and confocal images of the spine membrane (grey) depicting representative examples of macular or perforated assemblies and such consisting of 2 or 3 segments. **(B)** Percentage of macular, perforated, segmented 2, and segmented ≥ 3 PSD95 assemblies per spine for up to 120 min after cLTP induction (right) or control without stimulation (left). **(C)** The PSD95 coverage (covg) ratio was calculated by dividing the area covered with PSD95 (red) by the greatest extent of the synapse (dashed white ellipse). **(D)** PSD95 coverage ratio for control and cLTP potentiated spines over a 120 min time course (One-way ANOVA and Dunnett’s multiple comparison test; potentiated: -15 vs. 60 min: ***p = 0.0002, -15 vs. 120 min: ****p < 0.0001). Data represent mean ± SEM. Number of analysed PSD95 assemblies at -15, -5, 30, 60, 120 min: B: control: 227, 120, 221, 213, 92; potentiated: 110, 45, 102, 109, 60; D: control: 227, 120, 221, 213, 92; potentiated: 109, 45, 102, 109, 60.

To assess the reorganization of PSD95 quantitatively, we computed the coverage ratio of PSD95 in the synapse, which represents a measure for the degree of reorganization. To calculate the coverage ratio, we estimated the extent of the synapse by encircling all PSD95 of a synapse with an ellipse. Then we divided the area covered with PSD95 by the area of this ellipse (Figure 3C). Thus, the more black pixels there are between PSD95 segments or the larger a perforation is, the lower the coverage ratio. Analysing the coverage ratio for the whole time-course (Figure 3D, S3) showed that it was essentially constant for control spines; its average value varied between 0.782 ± 0.007 before cLTP at -15 min, 0.762 ± 0.008 at 60 min and 0.769 ± 0.013 at 120 min after cLTP. The coverage ratio in potentiated spines exhibited a significant decrease from 0.805 ± 0.012 before cLTP to 0.729 ± 0.013 at 60 min and to 0.705 ± 0.015 at 120 min after cLTP induction (Figure 3D). This supports the notion that the size expansion of PSD95 at 60 min after cLTP (Figure 2G) is accompanied by a remodeling of the PSD95 nano-organization.

### Synaptic GluA2 containing AMPAr area increases in size after cLTP

Next, we examined whether the activity-driven plasticity of PSD95 would also affect the nano-organization of AMPAr. An increase in the number of functional postsynaptic AMPAr is regarded as a fundamental mechanism of LTP^20^. AMPAr consist of tetramers of four different subunits - GluA1–GluA4 - which can be labeled specifically. In hippocampal neurons, most AMPAr contain the GluA2, which we therefore chose to label^21^. Since there is no live-cell-compatible marker for GluA2 we employed immunohistochemistry to label GluA2 in fixed, cultured neurons at different time points after cLTP induction. At the same time, we immunolabeled PSD95 and tagged the F-actin cytoskeleton by fluorescently-labeled phalloidin (Figure 4A). cLTP was induced for 5 min and thereafter the cells were transferred to standard ACSF and fixed either directly (0 min) or after 30, 60, or 120 min. Control cells were transferred at the same time points between ACSF solutions and fixed at the same time points. With a home-built super-resolution microscope, the immunolabeled PSD95 and GluA2 were recorded in super-resolution STED and actin was detected in confocal mode (Figure 4B). The size of the GluA2 and PSD95 nanostructures was assessed by encircling the outer extent of the nano-pattern (Figure S4A). We only considered synaptic GluA2, i.e., we only included GluA2 in the spine head, which touches or overlaps with the PSD95 nano-pattern. The areas of separate cluster in a spine head were summed to obtain the PSD95 and synaptic GluA2 area per spine head. After cLTP induction the area of synaptic GluA2 per spine head increased significantly from 0.056 (median; 0.046/0.072 lower/upper 95 % CI) µm^2^ at 0 min to 0.079 (0.064/0.098) µm^2^ after 30 min, to 0.098 (0.078/0.111) µm^2^ after 60 min and to 0.150 (0.129/0.179) µm^2^ after 120 min (Figure 4D, S4E). At the same time a significant increase in area was observed for PSD95 after 30 min, 60 min and 120 min (Figure 4C, S4D); for example, after 60 min the PSD95 assembly size increased from 0.169 (median; 0.154/0.181 lower/upper 95 % CI) µm^2^ for control to 0.249 (0.228/0267) µm^2^ after cLTP induction. This increase in size shifts and widens the frequency distribution (Figure S4D, E) similar to that of the mEPSC amplitudes (Figure 2C). The median size of GluA2 assemblies is on average 30 % of the PSD95 assembly size in control samples and 50 % of the PSD95 assembly size upon cLTP, i.e. they do not cover the entire PSD. Plotting the sizes of PSD95 assemblies vs. synaptic GluA2 assemblies revealed a moderate to strong correlation with a Pearson’s correlation coefficient *r* of 0.6-0.7 for all time points after cLTP induction (Figure 4F). Interestingly, the linear regression line of PSD95 area vs. synaptic GluA2 area was always relatively similar between cLTP and control (Figure 4F), indicating that PSD95 and synaptic GluA2 areas grow by a similar factor. As the size increase in the two variables after cLTP induction occurs at the same time scale in a correlated fashion, the increase in size of the PSD95 nanostructure is always accompanied by an increase in GluA2 nanostructure of the same factor for up to 2 h after cLTP induction. Remarkably, the regression line has a relatively large positive y-intersect at all time points (Figure 4F). This indicates that tiny GluA2 nanoclusters appear on PSD95 assemblies that are ∼ two times larger in size than the GluA2 nanocluster, whereas large GluA2 nanoclusters, for example with a size of 0.5 µm^2^, appear on PSD95 assemblies of a similar size on average.

**Figure 4.**
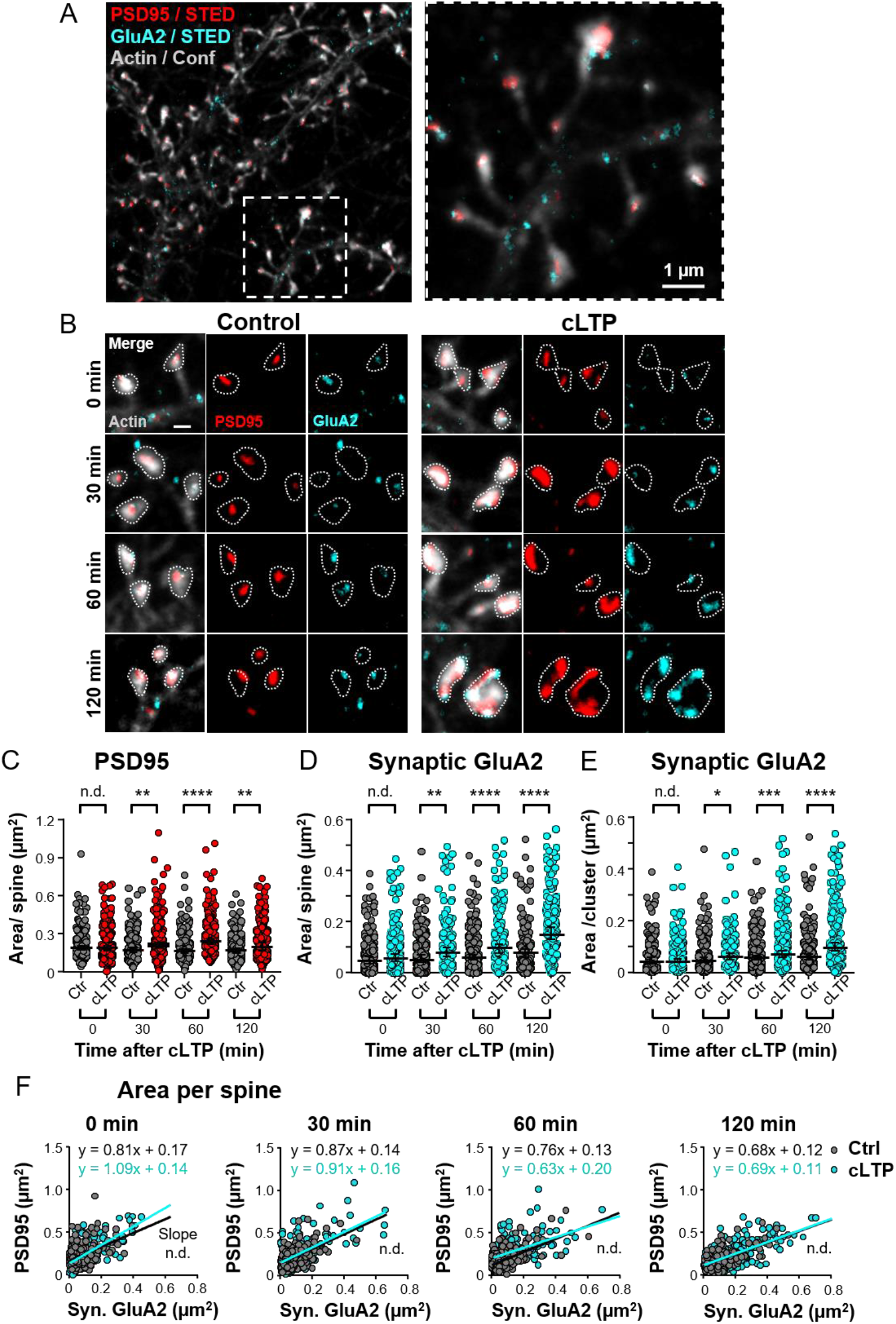
AMPAr nanocluster containing the GluA2 subunit and PSD95 nanostructures increase similarly in size after cLTP induction. **(A)** Two-color STED image of PSD95 and GluA2 (immunolabelling), and confocal image of actin (labelled with phalloidin) in hippocampal neuronal cell culture at 17 DIV. (**B)** Time series of hippocampal neuronal cultures fixed at 0, 30 min, 60 min, and 120 min after cLTP induction (right) or without stimulation (control, left). Scale bar: 500 nm. **(C)** Median ± 95 % confidence interval (CI) of PSD95 area per spine head (Mann-Whitney (M-W) test: 30 min: **p = 0.0027; 60 min: ****p < 0.0001; 120 min **p = 0.0020). **(D)** Median ± 95 % CI of total GluA2 area per synapse following cLTP induction and of control (M-W test: 30min: **p = 0.0018; 60 and 120 min: ****p < 0.0001). **(E)** Area of single synaptic GluA2 nanocluster with and without cLTP induction (median ± 95 % CI; M-W test: 30 min **p = 0.040; 60 min ***p = 0.001 and 120 min ****p < 0.0001). **(F)** Correlation between the size of the PSD95 and synaptic GluA2 area at different time points after cLTP induction or control samples fixed at the same time points; line shows linear regression. Pearson’s correlation coefficient r for control/cLTP: 0 min: 0.47/0.68; 30 min: 0.60/0.64; 60 min: 0.61/0.57; 120 min: 0.61/0.68. No significant difference in slope. Number of analysed spines (C–F): Control: 0 min: 188, 30 min: 199, 60 min: 222, 120 min: 188; cLTP: 0 min: 207, 30 min: 181, 60 min: 220, 120 min: 213.

Next, we tested whether the increase of the synaptic GluA2 area was due to an enlargement of existing clusters or the appearance of additional GluA2 nanoclusters at the synapse. Synaptic GluA2 containing nanoclusters increased significantly in size after 30 min and continued to increase after 60 and 120 min after stimulation (Figure 4E). However, we also found a small increase in the number of GluA2 containing nanoclusters per synapse, which was significant at 0 and 120 min after cLTP induction (Figure S4C). Thus, we observed mainly an increase in the GluA2 nanocluster size, which might be accompanied by an increase in the number of clusters per synapse. Of note, not all spine heads did contain GluA2. Spines without AMPAr at the synapse cannot be activated and are therefore called silent synapses. Supporting previous findings^11,22^, we observed that the number of silent synapses dropped over our time course; 120 min after cLTP induction only 1.4 % of the synapses were silent compared to 6.9 % without cLTP (Figure S4F). This corroborates the notion that LTP promotes the gradual insertion of new GluA2 at silent synapses.

### Collective pre- and postsynaptic enlargement

Recently, it was shown that pre- and postsynaptic elements align and form so-called molecular nanocolumns^9,10^. Therefore, the activity-induced enlargement and remodeling of PSD95 assemblies and AMPAr nanoclusters might be accompanied by corresponding presynaptic changes. Thus, we tested whether the size and nano-pattern of the presynaptic active zone protein Bassoon follows the same tendency as the PSD95 organization after cLTP induction. We fixed cultured hippocampal neurons at different time-points (0, 30, 60, and 120 min) after 5 min of cLTP induction (Figure 5B) as described above. We immunolabeled the neurons with antibodies against Bassoon and PSD95, and the F-actin cytoskeleton with phalloidin. Super-resolution imaging of Bassoon and PSD95 with STED microscopy resolved their synaptic nano-pattern and indicated an increase in size following cLTP (Figure 5A, B). The area covered by Bassoon and PSD95 was analysed by encircling the assemblies in a manner analogous to the analysis of GluA2 clusters. Only Bassoon assemblies that either overlapped with PSD95 or were directly opposite and thus most likely part of a synaptic contact were considered. After cLTP induction, Bassoon assemblies changed in size from 0.218 (median; 0.197/0.238; lower/upper 95 % CI) µm^2^ directly after cLTP (0 min) to 0.168 (0.155/0.177) µm^2^ after 30 min, increased up to 0.238 (0.213/0.256) µm^2^ after 60 min, and to 0.236 (0.212/0.259) µm^2^ after 120 min. While the first two time points were not significantly different from the control group, the size of Bassoon assemblies increased significantly 60 min and 120 min after cLTP (Figure 5D). PSD95 assemblies also increased in size after 60 and 120 min (Figure 5C); for example, after 120 min they increased from 0.174 (0.161/0.190) µm^2^ for control groups to 0.244 (0.225/0.270) µm^2^ after cLTP. Thus, the average size of Bassoon and PSD95 assemblies was very similar. The linear regression line of PSD95 area vs. Bassoon area was similar at 0, 30, and 60 min after cLTP and in control groups (Figure 5E); consequently, Bassoon and PSD95 assembly areas increase after cLTP induction on average by the same factor. However, the PSD95 assembly area increased slightly more than that of Bassoon after 120 min, which is indicated by a slightly larger slope of the regression line (Figure 5E). As such, the structural changes seen with the pre- and postsynaptic scaffolding proteins Bassoon and PSD95 upon cLTP are largely correlated during the remodeling for up to 120 min after cLTP induction.

**Figure 5.**
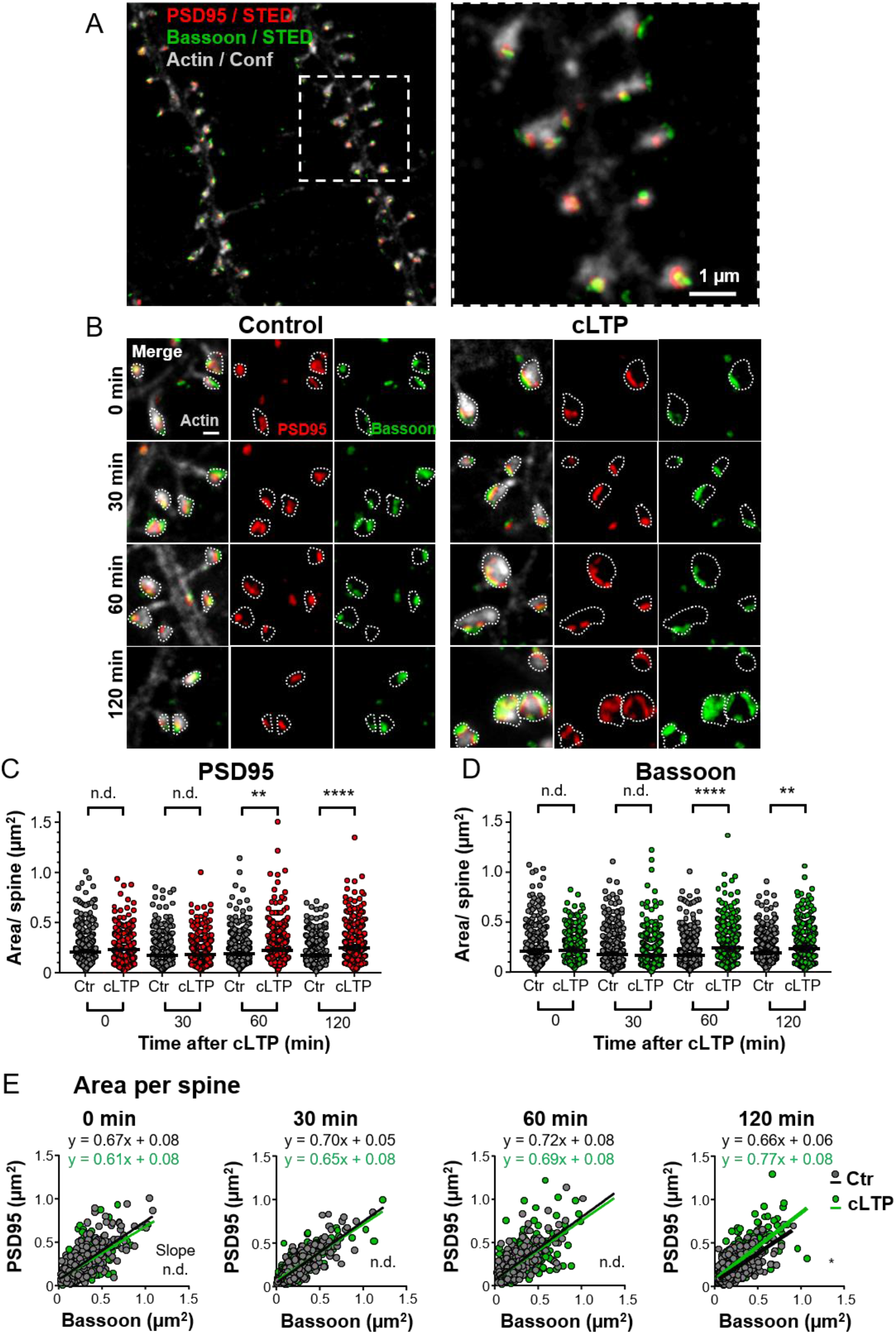
Coordinated increase of PSD95 and Bassoon assembly size after cLTP induction. **(A)** Two-color STED microscopy of PSD95 and Bassoon (immunohistochemistry labelling), and confocal image of actin (phalloidin labelling) in a hippocampal neuronal culture at 17 DIV. (**B)** Time series of neurons fixed at 0, 30 min, 60 min, and 120 min after cLTP induction (right) or control samples fixed at the same time points without stimulation (left). Scale bar: 500 nm. **(C)** Total PSD95 area per spine at 0, 30 min, 60 min, and 120 min after cLTP induction compared to control (median ± 95 % CI; M-W test: 60 min: **p = 0.0017; 120 min: ****p < 0.0001). **(D)** Same as (C), but for total Bassoon area facing PSD95 (M-W test: 60 min: ****p < 0.0001 and 120 min: **p = 0.0016). **(E)** Correlation between total PSD95 and total Bassoon area per spine at 0, 30 min, 60 min, and 120 min following cLTP compared to control; line shows linear regression. Pearson’s correlation coefficient r for control/cLTP: 0 min: 0.75/0.58; 30 min: 0.83/0.79; 60 min: 0.74/0.67; 120 min: 0.68/0.71. No significant difference in slope at 0, 30, 60 min; at 120 min: *p = 0.024. Number of analysed spines (C–E): Control: 0 min: 359, 30 min: 381, 60 min: 421, 120 min: 466; cLTP: 0 min: 340, 30 min: 310, 60 min: 424, 120 min: 370.

### Synaptic nano-organization of PSD95, GluA2, and Bassoon and its plasticity

Our time-lapse imaging of PSD95 in living, hippocampal slice cultures revealed a reorganization associated with an increase in the complexity of the PSD95 nano-pattern after cLTP induction (Figure 3B). Similarly, we also observed an increase in perforated and segmented PSD95 assemblies after cLTP induction in fixed, immunolabeled, cultured neurons (Figure S5B). As described for the time-lapse experiment, we categorized the PSD95 nano-pattern into macular, perforated, segmented 2, and segmented ≥ 3. At 120 min after cLTP induction, the distribution of the PSD95 nano-organization was significantly different from the control condition (Figure S5B). At this time point, the proportion of macular PSD95 decreased to 59 % compared to 82 % in control, the percentage of perforated PSD95 increased to 22 % versus 3 % in control, and the percentage of segmented 2 also increased to 18 % versus 12 % in control. Thus, the structural change of PSD95 assemblies from macular to segmented and/or perforated occurs with a delay after cLTP induction and was found in both organotypic hippocampal slice and dissociated hippocampal neuron culture.

We next examined whether these changes in the postsynaptic nano-organization are accompanied by corresponding presynaptic alterations and/or a reorganization of the GluA2-containing AMPAr clusters. We analysed the nanostructure of GluA2 and Bassoon assemblies in association with PSD95 after cLTP induction (Figure 6, S5A). The GluA2 nano-organization, an important functional indicator of cLTP, was much less complex than that of PSD95; we did not observe perforations, and therefore only assigned cluster numbers for GluA2 nano-organizations (Figure 6A). Figure 6B shows the frequency of the number of GluA2-containing nanoclusters as a function of the morphology of PSD95 assemblies. This plot reveals, for example, that macular PSD95 mostly contain only a single GluA2 nanocluster. Perforated PSD95 nanostructures, on the other hand, contain with similar frequency one, two, or three GluA2 nanocluster. Almost half of the segmented 2 PSD95 assemblies were occupied only with one GluA2 nanocluster, indicating that only one of the two PSD95 segments contained a GluA2 receptor patch. Similarly, ∼50 % of the segmented 3 PSD95 contained only one or two GluA2 nanocluster, and thus not each segment contained GluA2. Interestingly, silent synapses without GluA2 occurred almost exclusively on macular PSD95.

**Figure 6.**
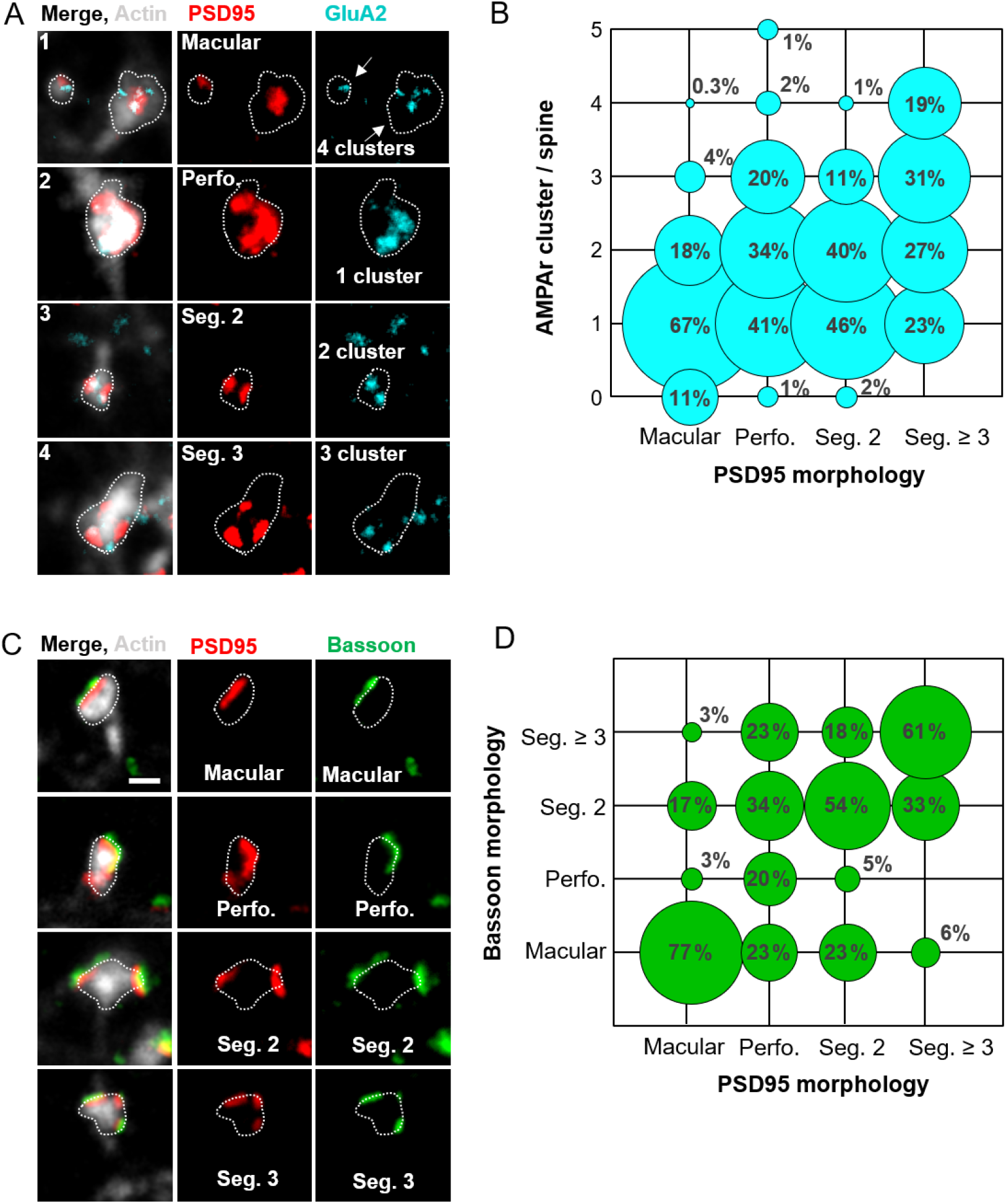
Similar nanoarchitecture across the synaptic scaffolds and glutamate receptors. **(A)** Categorization of PSD95 nano-organization (red) and GluA2 containing AMPAr nanocluster (blue). PSD95 structures are categorized into macular or perforated, or assemblies which consist of 2 or 3 separated segments; number of GluA2 nanoclusters located on PSD95 per spine. For an overview image refer to Figure S5A. Scale bar: 500 nm. **(B)** Frequency of the number of AMPA receptor clusters as function of PSD95 morphologies. **(C)** Examples of the different PSD95 morphologies and presynaptic Bassoon nanostructures. Scale bar: 500 nm. **(D)** Frequency of Bassoon morphologies on different PSD95 nano-organizations. Number of analysed spines (B) are the same as in Figure 4 and for (D) the same as in Figure 5; cLTP and control data were pooled for this structural correlation analysis.

The nanostructure of Bassoon can also be classified into macular, perforated, segmented 2, and segmented ≥ 3 (Figure 6C), and we found a strong similarity to the PSD95 nanostructure (Figure 6D). For example, macular Bassoon occurred together with macular PSD95 assemblies in 77 % of the cases; segmented 2 Bassoon appeared together with segmented 2 PSD95 and segmented 3 Bassoon with segmented 3 PSD95 in > 50 % of all cases. Although this conformity did not apply to perforated PSD95, all other Bassoon nanostructures occurred with similar frequency. Overall, these data document a high correlation and similarity between pre- and postsynaptic scaffolding proteins as regards their structural organization at the nanoscale.

## Discussion

In this study we used super-resolution time-lapse STED microscopy to explore activity-dependent synaptic plasticity based on changes in the size and nanostructure of assemblies of three major building blocks of glutamatergic synapses; the pre- and postsynaptic scaffolding proteins, Bassoon and PSD95, and the ionotropic glutamate receptor, AMPAr. All three protein assemblies increased slowly in size and complexity over the whole imaging period of two hours after cLTP induction; this enhancement was much slower than the rapid increase in synaptic strength as measured in mEPSCs or the spine head increase. The strong correlation between the shape changes of the pre- and postsynaptic scaffold protein assemblies and their temporal progression triggered by synaptic activity implies a subtle tuning between these structural parameters.

In the past, many studies showed a tight correlation between the size of the spine volume and the PSD^23^. Accordingly, the two parameters are often regarded as equivalent landmark indicators of changes in synaptic strength. In our measurements, however, the spine head increased already within 10 min of cLTP induction, while the increase in PSD95 assembly size was much slower, peaking at ∼60 min after cLTP induction. This is in line with electron microscopy^2^ and super-resolution microscopy^7,8^ studies reporting a temporal delay of the PSD95 assembly size increase after stimulation. This temporal decoupling may also explain why we found only a weak correlation between changes in PSD95 assembly and spine head size in mice after enhanced activity^6^. However, a temporal decoupling of this size correlation does not argue against a correlation at equilibrium or after adaptation. Interestingly, the relative increase in PSD95 assembly size was less than that of the spine head, indicating that cLTP has different dependencies to changes in PSD95 assembly and spine head. In essence, the essential characteristics of LTP formation cannot be explained only by changes in the size of PSD95 and spine head.

Our super-resolution imaging of endogenous PSD95 corroborates the complex shape of PSD95 assemblies we often observed before, also *in vivo*^5,6^. This shape is very diverse, so a simple classification into clusters, as often done in the context of super-resolution microscopy^3,11^, may understate their intricate function. We found that the complexity of PSD95 assemblies increased markedly after cLTP stimulation, peaking at the end of our measurement series of 2 h; for example, ∼18 % of the macular PSD95 assemblies transformed into a more complex shape with a perforation or split into two or more segments 2 h after cLTP induction (Figure 3B). We detected a complex PSD95 assembly shape less frequently than a previous study^2^. This could be due to the fact that our super-resolution imaging in 2D cannot resolve the nanostructure along the z-axis, which may underestimate the complexity of structures.

As previous studies indicated a strong structural and molecular coordination across the synapse in so-called trans-synaptic nanocolumns^9^ we performed a size and shape analysis of presynaptic Bassoon after cLTP induction. Assemblies of presynaptic Bassoon and postsynaptic PSD95 were very similar in size and shape; the size increase of Bassoon assemblies peaked at ∼ 60 min after stimulation, i.e. at a similar time scale as PSD95 assemblies. As such, the structural correlation between PSD95 and Bassoon assemblies was even stronger than proposed previously^9^. Thus, our data support the notion that synaptic activity can modulate a trans-synaptic structural correlation and alignment of molecular components. Intensive efforts are underway to determine how these structures align with each other across the synaptic cleft^24,25^.

A hallmark of LTP is the rapid potentiation of AMPAr-mediated postsynaptic currents and spine enlargement, which occur within a few minutes or less^26,27^. Current hypotheses pose that AMPAr are highly dynamic within the plasma membrane around the synapse and are trapped at synapses^20^. A remaining question is whether solely an increase in postsynaptic receptor number or also changes in the nanoscale organization of receptors contribute to changes in synaptic efficacy^28^. In particular, the low affinity of AMPAr for glutamate requires a tight alignment between AMPA receptors and glutamate release sites^3,25^, emphasizing a potential role of the synaptic nanoarchitecture. Previous studies indicate that AMPAr indeed align with presynaptic release sites^9^ and that AMPAr cluster more at the periphery of the PSD while NMDA receptors concentrate more at its center^29^. To investigate now activity-dependent changes of AMPAr, we super-resolved the nano-organization of GluA2 at different time points after cLTP induction. In line with the current literature, we found a clustered distribution of GluA2 within the synapse. Specifically, for GluA2 we did not observe complex structures such as perforations, indicating a different nano-architecture for GluA2 assemblies as compared to for PSD95 or Bassoon assemblies. The median size of the GluA2 assemblies was about 1/3 of that of PSD95 assemblies, i.e. synaptic GluA2 was restricted to a sub-region of PSD95 assemblies. Thus, the trapping of receptors by PSD95 does not occur at stochastically random PSD95 molecules, but involves an additional, yet unknown mechanism that causes cluster formation in subdomains within PSD95 assemblies. After cLTP induction, the synaptic GluA2 nanoclusters increased highly significantly in size and slightly in number, which deviates from a previously described pure modular architecture^10,11^. The increase in area size of synaptic GluA2 continued for up to two hours after cLTP induction and was therefore much slower than the reported fast increase of AMPAr mediated currents after glutamate uncaging^26,27^ and also slower than the increase of the mEPSC amplitudes that we observed after cLTP induction (Figure 2B). Therefore, our data do not indicate that the initial, fast increase of mEPSC amplitudes is due to an increase in synaptic GluA2 levels. The scaled increase of the mEPSC amplitude (Figure 2C, Figure S2B) and the increase and widening of the size distribution of GluA2 and PSD95 assemblies (Figure S4D, E) indicates that the synaptic strengthening involves most synapses and not just a few specific synapses. The strong, positive y-intersect of the linear regression line in Figure 4F, fitting the plot of PSD95 assembly sizes versus synaptic GluA2, indicates that the size ratio between PSD95 and GluA2 assemblies is different between small and large PSD95 assemblies. Thus, larger synapses with larger PSD95 assemblies harbor proportionally more GluA2. This fits with the hypothesis that larger synapses may have experienced synaptic strengthening and therefore integrated more AMPAr^20,30^.

Currently, the most convincing model for the formation of synaptic nanostructures is that of liquid-liquid phase separation^31^. Indeed, reconstituted PSDs were shown to self-organize into a web-like structure similar to native perforated PSD^31,32^. Moreover, CaMKII is able to drive the segregation of AMPA and NMDA receptors into separate nanodomains within the PSD^33^. Although these models so far include only a small selection of PSD proteins, they may adequately describe the formation or synaptic nanostructures.

## Supplemental figures

**Supplementary Figure 1:**
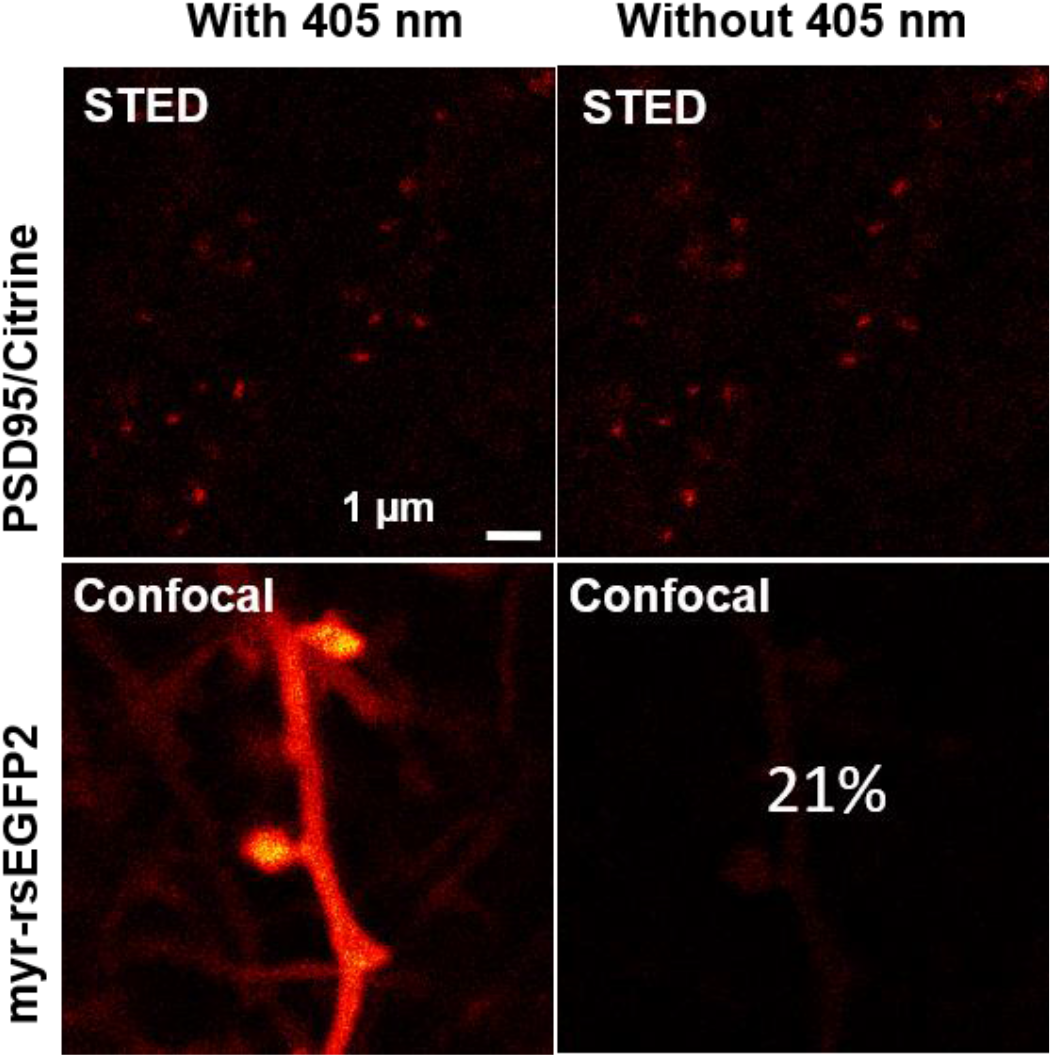
Imaging scheme. Neurons expressing either PSD95.FingR-Citrine (above) or myr-rsEGFP2 (below) in hippocampal organotypic slices. While both labels, Citrine and rsEGFP2, are excited at 480 nm, the latter can only be read out when it is switched to the on state by UV light (raw data); switching contrast between on and off state at the applied laser powers is 21 %.

**Supplementary Figure 2:**
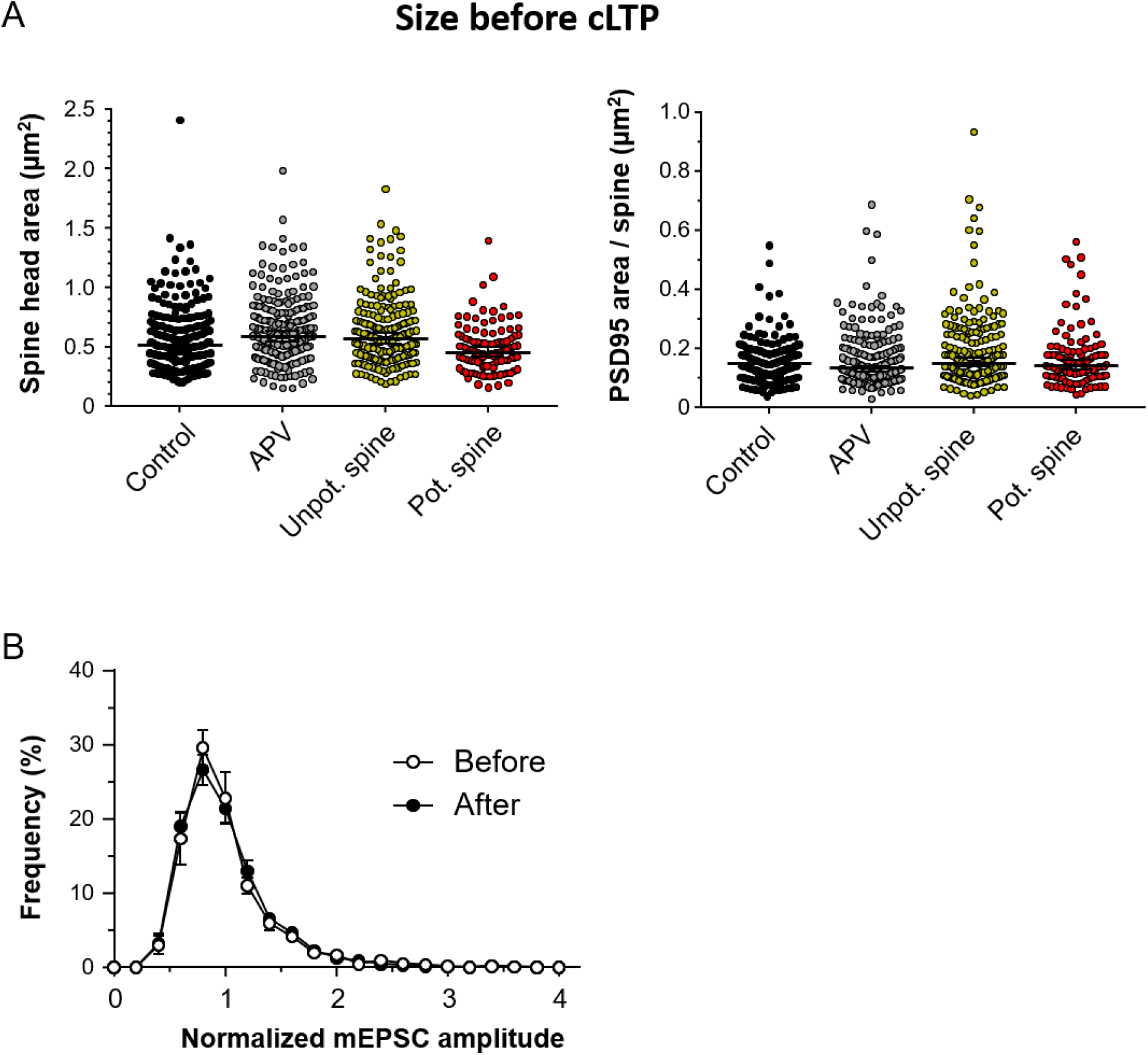
**(A)** Size distribution of spine heads (left) and PSD95 area (right) per spine head before treatment. Mean size of spines heads (lower/upper 95 % CI): Control: 0.57 (0.53/0.61) µm^2^; APV: 0.63 (0.60/0.67) µm^2^; Unpotentiated spine heads: 0.62 (0.58/0.66) µm^2^; potentiated spine heads: 0.48 (0.45/0.52) µm^2^. Mean size of PSD95 assemblies (lower/upper 95 % CI): Control: 0.15 (0.14/0.16) µm^2^; APV: 0.16 (0.15/0.17) µm^2^; PSD95 on unpotentiated spines: 0.19 (0.17/0.20) µm^2^; PSD95 on potentiated spines: 0.17 (0.15/0.19) µm^2^. **(B)** Frequency distributions of normalized mEPSC amplitudes before (open circle, -15 min) and after cLTP induction (closed circle, 65 min) are very similar in shape. Same dataset as Figure 2C.

**Supplementary Figure 3:**
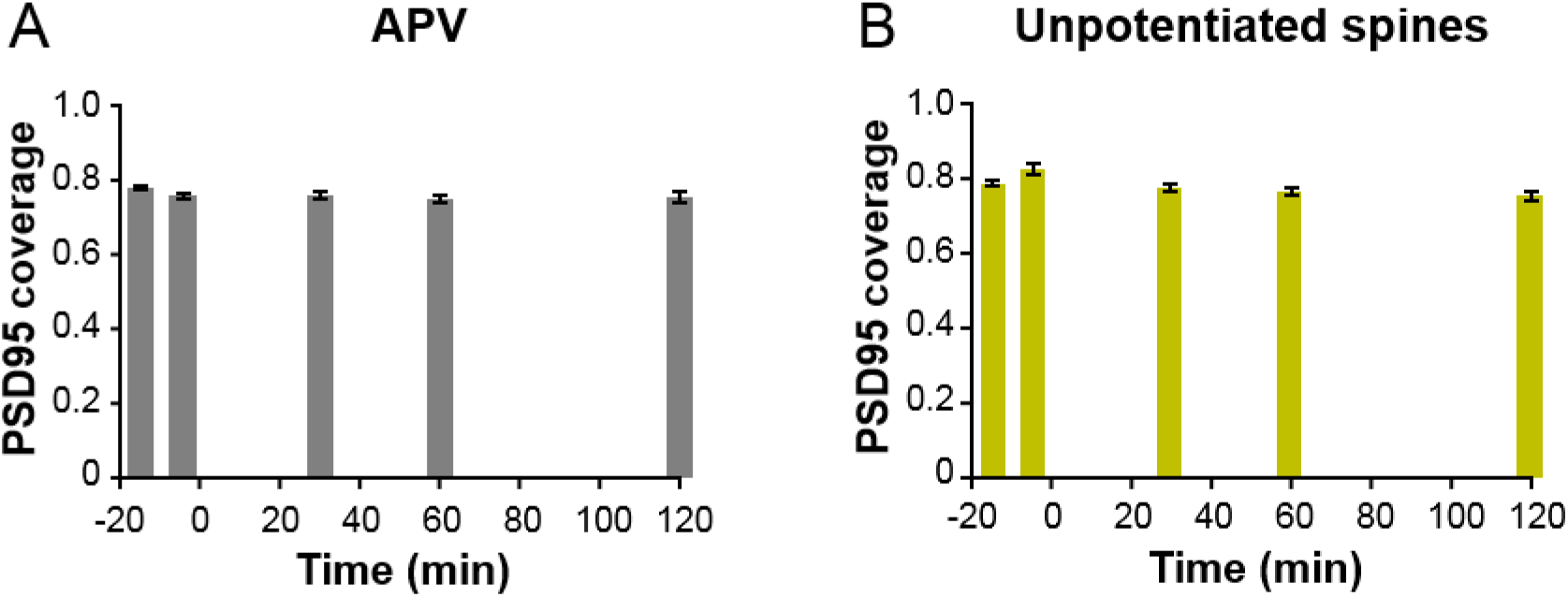
PSD95 coverage ratio for APV treated and unpotentiated spines over a 120 min time course (One-way ANOVA and Dunnett’s multiple comparison test; no significant changes over time).

**Supplementary Figure 4:**
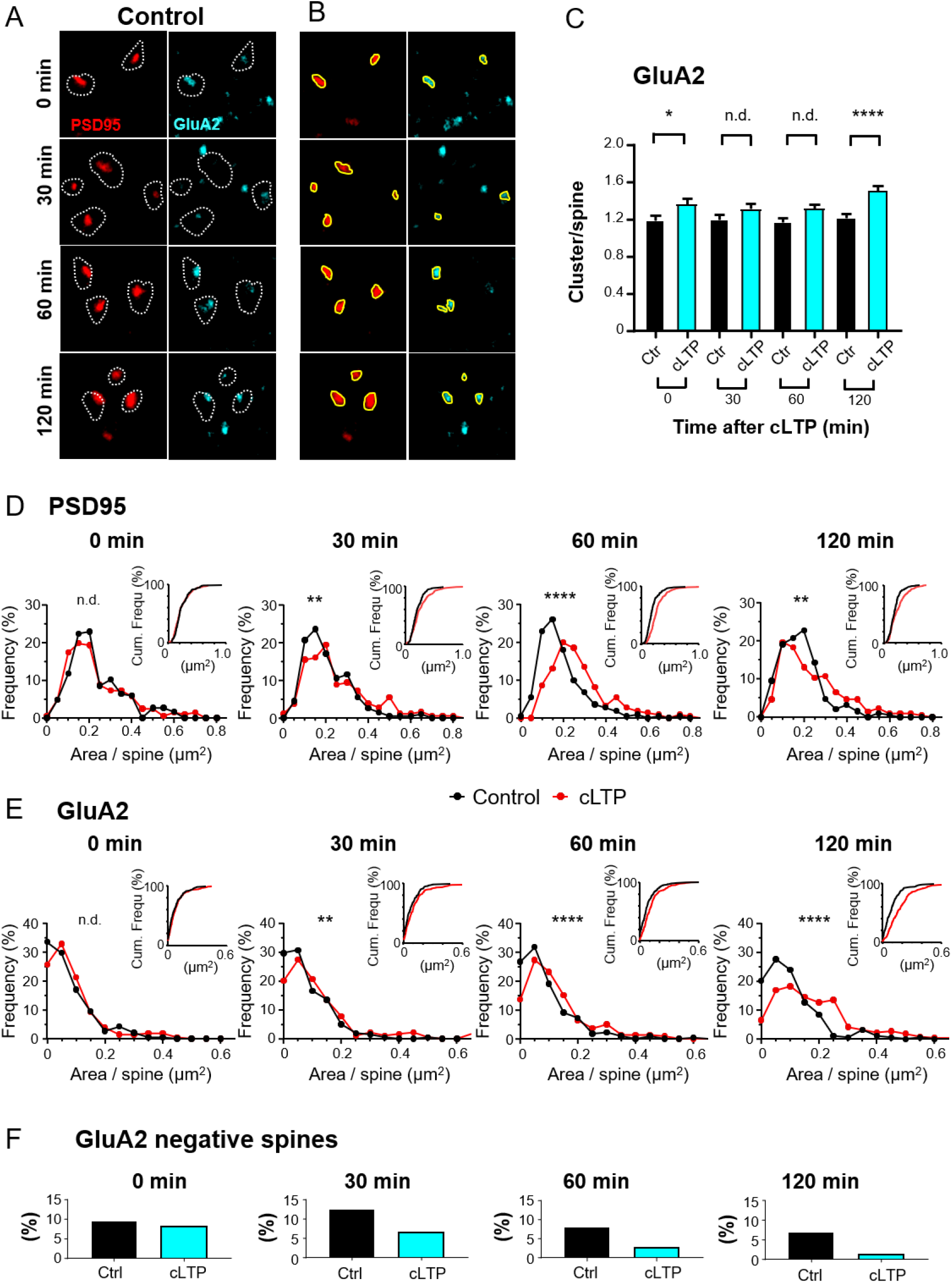
**(A, B)** Size analysis of PSD95 assemblies and GluA2 containing AMPAr cluster exemplified on the control group of Figure 4; all assemblies per spine were outlined (B) to calculate the area covered. **(C)** The number of AMPA receptor (AMPAr) clusters per PSD95 nano-organization after cLTP compared to control; data shown as mean ± SEM (M-W test: 0 min *p = 0.039 and 120 min ****p < 0.0001). **(D, E)** Frequency distribution and cumulative frequency (inset) of PSD95 area (D) and GluA2 area (E) per spine; data and test same as in Figure 4C, D. **(F)** Percentage of spines without AMPA receptor clusters based on PSD95 organizations following cLTP compared to control; 0 min, cLTP: 8.2 % vs Ctrl: 9.6 %; 30 min, cLTP: 6.6 % vs Ctrl: 12.6 %; 60 min, cLTP: 2.7 % vs Ctrl: 8.1 %; 120 min, cLTP: 1.4 % vs Ctrl: 6.9 %.

**Supplementary Figure 5:**
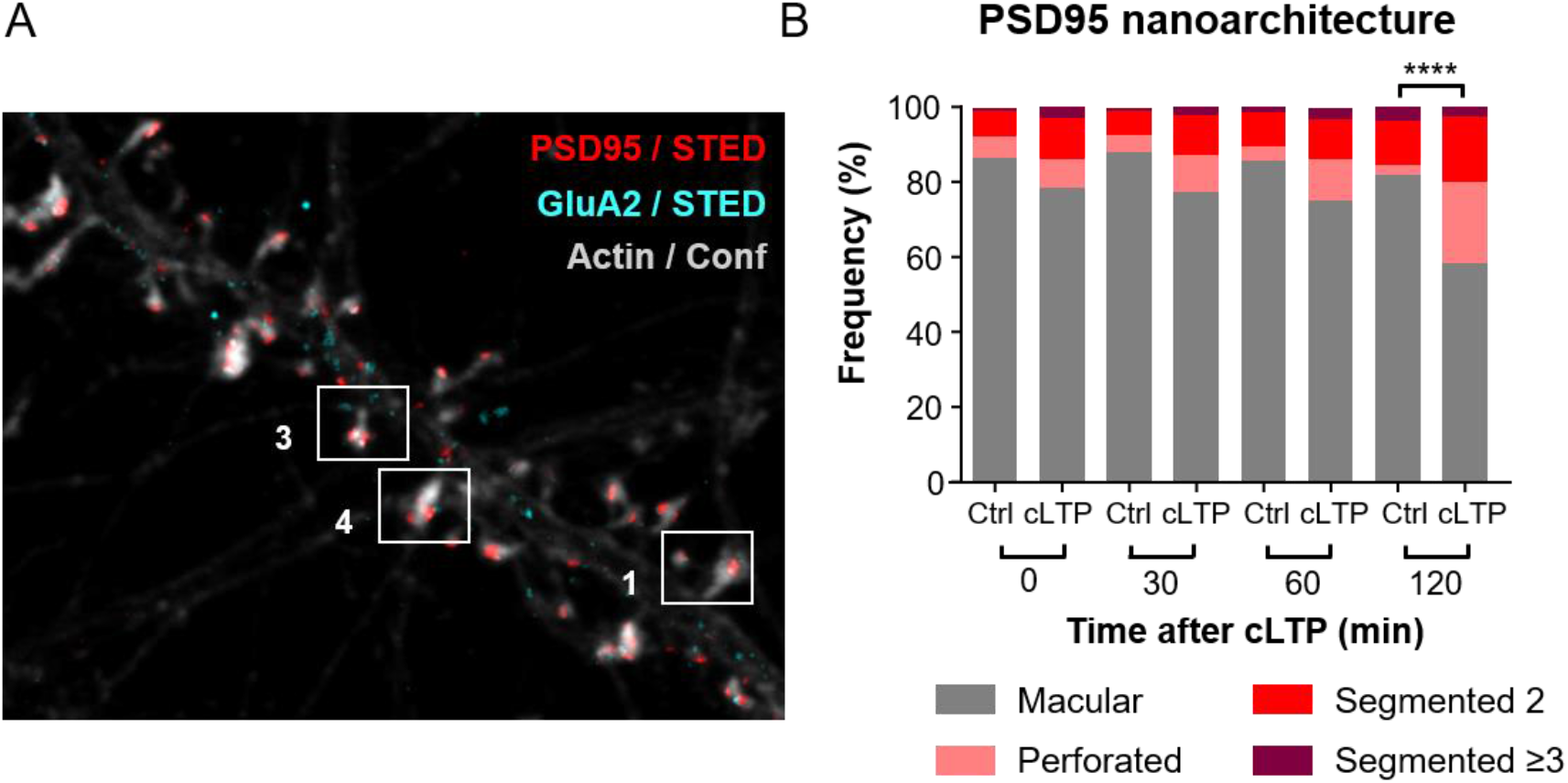
**(A)** Overview image of PSD95, GluA2 and actin labeled neuronal culture; boxed and numbered are spines shown in Figure 6. **(B)** Frequency of PSD95 morphologies at 0, 30 min, 60 min, and 120 min following cLTP compared to control (Ctrl); the distribution of morphologies is significantly different between cLTP and control after 120 min (Kolmogorov-Smirnov (K-S) test; ****p < 0.0001).

## Methods

### Neuronal hippocampal cultures

Primary neuronal hippocampal cell cultures were prepared according to^34^ from the hippocampi of Wistar rats of both sex at day P1. Cells were plated at density of 180,000 cells/ml on 18 mm round coverslips of thickness #1.5 (Marienfeld, # 0117580) in 12-well plates and maintained at 37°C with 5 % CO_2_.

### Organotypic hippocampal slices

Organotypic hippocampal slice cultures were prepared from P5 C57BL/6 wild-type mice of both sex according to^35^. In brief, mice were decapitated and the hippocampus was extracted and sliced into coronal sections of 300 µm thickness. The slices were placed on small pieces of Polytetrafluoroethylene (PTFE) membrane, pore size 0,45 µm (Millipore, # FHLC01300) and cultured on cell culture insert of 0.4 µm pore size (Millipore, # PICM03050) placed in a 6-well plate and then incubated at 37°C with 5 % CO_2_ for 18 to 23 days *in vitro* (DIV). The inhibitor mix containing Ara-C (Sigma-Aldrich, # C6645), Uridine (Sigma-Aldrich, # U3750) and 5-Fluoro-2’-desoxyuridine (Sigma # F0503), was added to a final concentration of 3 µM on the third day and the medium was exchanged three times per week.

### Live-cell labelling with rAAV

Plasmid cloning and production of recombinant adeno associated virus (rAAV) is described in^13^. The membrane of dendrites and spine are labeled by transduction with rAAV-hSyn-DIO-myr-rsEGFP2-LDLR(ct)-WPRE which encodes for the reversible photoswitchable (rs) green FP rsEGFP2^36^. The rsEGFP2 tag is attach to an N-terminal myristoylation motif (myr) that promoted membrane labelling, and the C-terminal (Ct) cytoplasmic domains of low density lipoprotein receptor (LDLR) to target the protein to the dendrite^37^. The expression is cre-dependend by insertion of a double-floxed inverted open reading frame (DIO) under the control of the neuron-specific human synapsin-1 promoter (hSyn).

Endogenous PSD95 is visualized by transducing rAAV-ZFN-hSyn-DIO-PSD95.FingR-Citrine-reg.-WPRE^6,13^ encoding for a transcriptionally regulated recombinant antibody-like-protein called FingR (Fibronectin intrabodies generated with mRNA display) fused to the yellow FP Citrine^14^.

Both rAAV were transduced together with a low concentration of the cre-recombinase encoding virus rAAV-hSyn-CRE-WPRE^18^ in the CA1 region of an organotypic hippocampal slices 2 days after preparation. The virus was injected via a pulled needle of a borosilicate glass capillary (ID: 0.68mm, OD: 1.2mm; Kwik-fill, World Precision Instruments Inc., # 1B150F-4) that was angled at 50° to the slice using a stereotactic micromanipulator (SM-11, (Narishige Scientific Instrument Lab.). ∼50 nl of the virus mixture was pressure injected with ∼10 pulses at 15 psi via an Intracellular Microinjection Dispense System (PICOSPRITZER III, Parker Instrumentation).

### Chemical LTP

For immunostaining experiments, hippocampal cultured neurons were used between 16 to 21 DIV. To induce cLTP, we treated the neurons for 5 minutes at 37°C with modified artificial cerebrospinal fluid (ACSF) in which magnesium was removed, and 200 μM glycine (Sigma-Aldrich, # G8790) and 20 μM bicuculline (Hello Bio, # HB0893) were added instead^38^. The composition of ACSF for the hippocampal cultured neurons was as follows (mM); 2 MgCl_2_, 105 NaCl, 2,4 KCl, 10 HEPES, 10 D-glucose, 2 CaCl_2_ at pH 7.4, and ∼240 mOsm^40^. The treated neurons were immediately fixed (time point 0 min) or kept in ACSF and fixed after 30, 60, or 120 min post cLTP induction. As control, the hippocampal cultured neurons were incubated in ACSF and fixed at the same time points.

To record the cLTP with live-cell STED imaging of hippocampal organotypic slices, we also used the modified ACSF for 10 minutes. However, the composition of the ACSF used here is as follows (mM); 2 MgCl_2_, 128 NaCl, 2 KCl, 10 KH_2_PO_4_, 26 NaHCO_3_, 10 glucose, 2 CaCl_2_, pH 7.4 and ∼ 320 mOsm. Before induction of cLTP, the slices were maintained in ACSF solution supplemented with 50 µM D*-*2-amino-5-phosphonovaleric acid (APV, Hello Bio, # HB0225) for 20 minutes to block the activity mediated by NMDA receptors. After cLTP induction, the slices were perfused with ACSF for 2 h. For control, the solutions were changed at the same time points but all solutions contained ACSF supplemented with 50 µM APV for blocking (marked as APV throughout the manuscript). To investigate the basal activity, the hippocampal organotypic slices were maintained in the ACSF during the whole experiment (marked as CONTROL). During the live-STED imaging sessions, the hippocampal slices were preserved at 30°C via a heated platform (QE-2, Warner Instruments, LLC) and solution heater (SF-28, Warner Instruments, LLC); both are controlled by a dual temperature controller (TC-344C, Warner Instruments, LLC). The ACSF solution was continuously infused with 5 % CO_2_ and delivered with a flow of ∼1 ml/min (MINIPULS 3 Peristaltic Pump, Gilson).

### Immunocytochemistry

The cultured hippocampal neurons were fixed in a 3 % v/v glyoxal solution (Sigma-Aldrich, # 128465) containing 0.75 % acetic acid (Carl Roth, # 3738) for 1 hour according to^39^. The fixative was adjusted to pH 4 for staining Bassoon/PSD95 and to pH 5 for staining GluA2/PSD95. After fixation, cells were quenched in 0.1 M glycine in PBS and then permeabilized for 30 min in a blocking solution of 2 % normal goat serum (Sigma-Aldrich, # G9023) and 0.1 % Triton X-100 (Sigma-Aldrich, # T8787) in PBS. The primary antibodies rabbit anti-bassoon (Synaptic Systems, # 141013, dilution 1:500), mouse anti-PSD95 (Neuromab, # 75-028, dilution 1:300), and Alexa Fluor 488 phalloidin (Invitrogen, # A12379, 1:600) were diluted in blocking solution and the neurons were incubated for 2 h at room temperature.

To label AMPA receptors in the plasma membrane the N-terminal extracellular domain of GluA2 was stained with monoclonal mouse anti-GluA2 (Millipore, # MAB397, 1:500) for 2 hours without prior permeabilization. Thereafter, cells were permeabilized in blocking solution supplemented with 0.1 % Triton X-100 for further intracellular labeling of PSD95 and actin. The samples were incubated with rabbit anti-PSD95 (Cell signaling, # 3450, 1:300) and Alexa Fluor 488 phalloidin (1:600) diluted in blocking solution (with Triton X-100) for 2 h at room temperature.

After washing, the secondary antibodies anti-rabbit STAR RED (Abberior, # STRED-1002, dilution 1:50) and anti-mouse Alexa Fluor 594 (Thermo Fisher Scientific, # A-11005, dilution 1:100) were incubated at 4°C overnight in blocking solution. Finally, the coverslips were mounted with Mowiol (Carl Roth, # 0713).

### Live-cell STED imaging

For live-cell imaging of the organotypic hippocampal brain slices the STED microscope was configured as describe in^13^ with the following minor adaptions. The images were collected with a water dipping objective of 1.2 numerical aperture equipped with a correction collar (HC PL APO 63x/1,20 W CORR CS2, Leica Germany, # 506356). Fluorescence of both, rsEGFP2 and Citrine, was detected between 510–560 nm by a bandpass filter (AHF Analysentechnik). Images of PSD95 and the membrane label were recorded sequentially. Firstly, PSD95 was superresolved using the blue light excitation together with the STED beam; immediately afterwards, a confocal image of the myristoylation tag was recorded at the same dwell-time and excitation power, without the STED beam, but with additional UV light to switch the rsEGFP2 to the on state^13^.

The organotypic hippocampal slices were imaged at 18 to 23 DIV. Therefore, a slice was placed in a 35 mm petri dish and attached to the bottom of the dish with silicon glue (twinsil, picodent, # 1300 1000) to prevent the slice from floating away or moving during time-lapse imaging. The slice was continuously perfused with ACSF and heated as described above. Confocal and STED x,y-images were recorded in a stack over 2.5 µm at a distance Δz of 500 nm and with a dwell-time of 4 µs with the software Imspector (Abberior Instruments). The dimension of the x,y-image was 30 × 30 µm with a pixel size of 30 nm square. Fast coarse confocal overview images were recorded before each STED image stack to realign the field of view with the earlier recorded image area. For time-lapse imaging STED and confocal stacks were acquired 4 times in two different settings: before cLTP, during cLTP, 30 and 60 min after cLTP, or before cLTP, 30, 60, and 120 min after cLTP. The power of the blue excitation beam was 5.5 µW, that of the UV light for switching 2 mW and STED was performed with ∼15 mW measured at the entrance pupil of the objective, respectively.

A home-built inverted STED microscope described in^18^, was slightly modified to image the triple color immunostaining of Bassoon, PSD95 and actin or respectively GluA2. A blue laser line was added to excite Alexa Fluor 488 at 488 nm with 8 µW (Cobolt 06-MLD, HÜBNER Photonics); its fluorescence was detected after a confocal pinhole at 510–560 nm. STAR RED was excited with 18 µW red light of 630 nm central wavelength and detected at 692/40 nm. Alexa Fluor 594 was excited with 15 µW in the orange at 586 nm and detected at 620/14 nm. Both, STAR RED and Alexa Fluor 594 were depleted at 775 nm with 230 mW power measured at the entrance pupil of the objective. All images were acquired quasi-simultaneously by repeating each line three times with alternated excitation wavelengths and detection channels. Images were collected in stacks of 5 pictures of 30 × 30 μm in x, y over 2 µm at distances Δz of 400 nm; the x/y pixel size was 20 nm squared and pixel dwell-time 5 µs.

### Image processing

For the live-cell imaging experiments, the size of the spine head and corresponding PSD95 area were analyzed with Fiji/ImageJ^40^. The images were first smoothed to average each pixel with 3 × 3-pixel neighborhood and the background was subtracted. To measure the spine head and PSD95 area each was encircled with the freehand selection tool. The changes in both the PSD95 area (Δ PSD95 area) and the spine head area (Δ spine head area) at each time point during the time course of the experiment was calculated as ΔA/A_0_. The value A_0_ corresponds to the initial time point (before cLTP); ΔA refers to the difference between the areas at time point t (0, 30, 60 or 120 min after cLTP) with the initial time point A_0_. To normalize to control, the normalized control values were subtracted at each time point.

PSD95 nano-organizations were analyzed according to their shape. When the spine head contained only one PSD95 assembly without any perforation or nanostructure, it was assigned a macular morphology. PSD95 with a U-shape, ring-like or more complex shape, which was continuously connected was assigned a perforated morphology; to account for noise, holes in the center or discontinuities were only considered if they were at least 3 × 3 pixels (90 nm x 90 nm) in size. PSD95 forming more than one cluster in a spine head which were separated by at least 3 pixels were marked segmented and the number of clusters were counted.

The spreading of PSD95 in a synapse was estimated by computing the coverage ratio. By that means, we estimated the extent of the synapse with an ellipse encircling the complex PSD95 assemblies. The coverage was computed by dividing the area covered with PSD95 by the area of the encircled ellipse, the outer expansion; thus 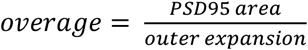. In this way the coverage drops the larger a hole in a perforation gets or the more two clusters are separated. The PSD95 area was obtained by encircling PSD95 with the freehand selection tool in FIJI; the area of the ellipse encircling the PSD95 assembly was obtained by fitting the length and the width of PSD95 assembly with an ellipse in Fiji.

The two-color STED images of immunolabeled PSD95 and GluA2 or Bassoon, respectively, were analyzed in Fiji as follows. The images were smoothed and the background was subtracted. The area was obtained by encircling each protein assembly manually. The size of segmented PSD95 in a spine head was summed. Bassoon was only included when it was contacting PSD95. The shape of the PSD95 and Bassoon nano-pattern was analyzed as described above and assigned a macular, perforated or segmented shape. GluA2 did not show perforations and thus only the number of clusters was counted; included were only clusters located on PSD95 nano-organizations, and clusters were required to be separated at least by 3 pixels (60 nm).

### Electrophysiological recording

Spontaneous miniature excitatory postsynaptic synaptic current (mEPSC) was recorded in CA1 pyramidal neurons of organotyptic hippocampal slice culture at DIV 16–21 using the whole cell patch recording mode under voltage-clamp conditions at -70mV, using a double patch-clamp amplifier (EPC-10, HEKA) with Patchmaster software; the recording electrode (2.5–3.5 MΩ) contained internal solution (mM) with 138 K-gluconate, 16.8 HEPES, 10 NaCl, 1 MgCl_2_, 4 ATP-Mg, 0.3 GTP-Na and 0.25 K-EGTA at pH 7.38 and 310 mOsm. The external solution was carbogen-saturated ACSF containing (mM) 120 NaCl, 20 KCl, 10 KH_2_PO_4_, 26 NaHCO_3_, 10 glucose, 2 CaCl_2_, with a pH of 7.4 and 302 mOsm. The AAV-PSD95-FingR-Citrine and AAV-myr-rsEGFP2 were overexpressed in the neurons by the viral system at 2 DIV. The organotypic hippocampal slices were constantly supplied with the ACSF before and during the recording. For cLTP, we record the mEPSC before and after treatment with modified ACSF (Mg^2+^ free, 200 μM glycine and 20 μM bicuculline) which was applied for 10 min. After treatment, mEPSCs were recorded for another hour in the standard ACSF. All extracellular solutions for mEPSC were continuously infused with tetrodotoxin (TTX) (1 µM, Tocris, # 1078) and 20 μM bicuculline.

### Statistical analysis

To compare more than 2 conditions, the unpaired one-way analysis of variance (ANOVA) was used for normally distributed data. Kruskal-Wallis (K-W) test was performed for non-normally distributed results. The comparison between two conditions was performed with the Mann-Whitney (M-W) test. Cumulative distributions were tested with the Kolmogorov-Smirnov (K-S) test. The respective test is listed in the figure legend. The number of experiments and analyzed structures are listed in the figure legend. All statistical analyses were generated via GraphPad Prism.

### Data availability

Source data files of all analyzed data will be provided.

## Acknowledgement

We thank Dr. Heba Ali, Max Planck Institute for Multidisciplinary Sciences, Göttingen, for help with analysis by providing macros in Fiji. This work was supported by the Göttingen Graduate Center for Neurosciences, Biophysics, and Molecular Biosciences (VCF) and by the Deutsche Forschungsgemeinschaft (DFG, German Research Foundation) within the DFG Research Center and Cluster of Excellence (EXC 171, Area A1) ‘Nanoscale Microscopy and Molecular Physiology of the Brain’ (WW, KIW) and the Collaborative Research Center 889 ‘Cellular Mechanisms of Sensory Processing’ (Project B7) (VCF, KIW).

## Author contributions

Conceptualization (VCF, KIW); cell culture, cLTP and imaging experiments (VCF); cloning and virus production (WW); electrophysiology (CKL, JSR, NB); analysis (VCF, KIW, CKL, JSR); coordination of the project (KIW); writing original draft (VCF, KIW); all authors revised and agreed on the manuscript.

